# Systematic Perturbation of an Artificial Neural Network: *A Step Towards Quantifying Causal Contributions in The Brain*

**DOI:** 10.1101/2021.11.04.467251

**Authors:** Kayson Fakhar, Claus C. Hilgetag

## Abstract

Lesion inference analysis is a fundamental approach for characterizing the causal contributions of neural elements to brain function. Historically, it has helped to localize specialized functions in the brain after brain damage, and it has gained new prominence through the arrival of modern optogenetic perturbation techniques that allow probing the functional contributions of neural circuit elements at unprecedented levels of detail.

While inferences drawn from brain lesions are conceptually powerful, they face methodological difficulties due to the brain’s complexity. Particularly, they are challenged to disentangle the functional contributions of individual neural elements because many elements may contribute to a particular function, and these elements may be interacting anatomically as well as functionally. Therefore, studies of real-world data, as in clinical lesion studies, are not suitable for establishing the reliability of lesion approaches due to an unknown, potentially complex ground truth. Instead, ground truth studies of well-characterized artificial systems are required.

Here, we systematically and exhaustively lesioned a small Artificial Neural Network (ANN) playing a classic arcade game. We determined the functional contributions of all nodes and links, contrasting results from single-element perturbations and perturbing multiple elements simultaneously. Moreover, we computed pairwise causal functional interactions between the network elements, and looked deeper into the system’s inner workings, proposing a mechanistic explanation for the effects of lesions.

We found that not every perturbation necessarily reveals causation, as lesioning elements, one at a time, produced biased results. By contrast, multi-site lesion analysis captured crucial details that were missed by single-site lesions. We conclude that even small and seemingly simple ANNs show surprising complexity that needs to be understood for deriving a causal picture of the system. In the context of rapidly evolving multivariate brain-mapping approaches and inference methods, we advocate using *in-silico* experiments and ground-truth models to verify fundamental assumptions, technical limitations, and the scope of possible interpretations of these methods.

**Author summary:** The motto *“No causation without manipulation”* is canonical to scientific endeavors. In particular, neuroscience seeks to find which brain elements are causally involved in cognition and behavior of interest by perturbing them. However, due to complex interactions among those elements, this goal has remained challenging.

In this paper, we used an Artificial Neural Network as a ground-truth model to compare the inferential capacities of lesioning the system one element at a time against sampling from the set of all possible combinations of lesions.

We argue for employing more exhaustive perturbation regimes since, as we show, lesioning one element at a time provides misleading results. We further advocate using simulated experiments and ground-truth models to verify the assumptions and limitations of brain-mapping methods.

## Introduction

One of the most challenging goals of neuroscience is to identify neural elements – brain regions, populations, neuronal circuits, and large-scale networks – that pivot cognition and behavior[1]. During the past two decades, brain mapping flourished with the help of neuroimaging techniques that associate elements and functions. Arguably though, the first method of mapping brain function, i.e., by studying lesions, yet has an authoritative role in establishing causation since it indicates the *necessity* of the element for a given function[2,3]. With this inferential capacity, though, comes practical and methodological difficulties that might deliver deceiving results [4,5]. Crucially, since the ground-truth causal processes in the brain are unknown, the limitations of how functional contributions are mapped to interacting neural elements are not fully resolved, and thus conventional lesion-based methods are left with unverified assumptions and unexplored alternatives[5].

From a practical point of view, the scale of available human lesion datasets is nowhere on a par with those used in and produced by correlative approaches. This is in particular problematic since, as it is shown, even by focusing on single local lesions, mass-univariate lesion analysis provides systematically biased maps while multivariate approaches require a considerable amount of data to remedy the problem [2,6]. Additionally, with invasive approaches and in animal models, the sheer number of elements in the brain makes it practically impossible to lesion all of them exhaustively in all but very small nervous systems[7,8].

Practical issues aside, cognitive functions emerge from interactions of distributed neural elements that make it challenging to isolate the functional contributions of individual units[5,9] while the established approach assumes to disassemble such coalitions by removing individual elements and assigning the resulting behavioral change as the elements’ contribution[3,10]. Historical cases of lesion inference after brain damage, in patients such as Phineas Gage and Henry Molaison (‘HM’) [11], as well as modern cutting-edge experimental tools employing opto- and chemogenetics that temporarily perturb the brain with astonishing spatiotemporal precision [12,13], mostly follow the same “Single-element Perturbation Analysis (SPA)” framework. It is important to note that a SPA study might have a multivariate approach by incorporating many variables, e.g., lesion volume, but one neural element is perturbed -or fed into a statistical model- at a time, whether the element is single neurons, a local circuit, or a brain region[5,14]. Put differently, neural elements produce behavior as spatially distributed, *interacting* coalitions[15–17] while the established methods mainly map the observed effects on local processes. Consequently, the SPA framework might overlook the subsequent effects that local lesions might have on the system as a whole[18]. Paradoxical lesion effects and, in particular, the “Sprague Effect” are intriguing phenomena to illustrate potential issues with this approach[19,20]. The Sprague effect describes a scenario in which disruptions in behavior caused by a first lesion revert to normal after a second lesion[20,21]. In other words, lesioning region *i* disrupts the behavior, providing apparently compelling evidence for its “necessity” for the behavior, while a subsequent lesion to another region *j* restores the behavior showing the redundancy or degeneracy of the contribution of *i*.

Different hypotheses have attempted to explain this unexpected result based on the inhibitory relationship between competing regions[22–24] or neuronal plasticity and the increased excitatory-to-inhibitory synaptic balance of the circuit[25]. Essentially, the Sprague effect points towards a more complex causal relationship in the brain rather than a single neural element-to-single function relationship, indicating how misleading it can be to assign functions to neural elements relying on individual lesions[18].

To further emphasize on this point, Jonas and Kording performed an exhaustive SPA of every transistor in a microprocessor to see if it reveals a meaningful causal picture of a system that we have confound-free access to, virtually, every computational unit of it[26]. They found a subset of transistors that perturbed, would disrupt the function of the microprocessor; however, they declared the results *“grossly misleading”* since *“The transistors are not specific to any one behavior [...] but rather implement simple functions”* [26]. Their results suggest that even by perturbing every relevant unit of a system, one at a time, we are still far from a coherent causal understanding of what is doing what and indeed prone to miss-attribute individual elements to a behavior that is emerged from complex interactions of many units.

In this work, we use an alternative approach known as “Multi-perturbation Shapley value Analysis (MSA)” that, in contrast to SPA, derives causal contributions of elements from permuting all combinations of multi-element lesions [27,28]. MSA is based on Shapley value, a game-theoretical metric that is used for fair distribution of costs, gains, or resources among players of a cooperative game[29]. In the context of neuroscience, players are arbitrarily defined neural elements that are ranked according to their contributions to an arbitrary quantified behavior or cognitive function[30,31].

Inspired by the provocative findings of Jonas and Kording and further investigating the inferential capabilities of SPA and MSA frameworks, we decided to use a ground-truth model and systematically perturb its components. Therefore, we perturbed all neurons and connections of a compact ANN, both one element at a time, that is, through SPA, or many combinations of elements, that is, by MSA. We used an ANN instead of a microprocessor to capture the whole spectrum of the behavioral performance instead of a binary state of disturbed versus functional performance. Moreover, to train the network we specifically used an evolutionary algorithm focused on the network’s topology to avoid handcrafting and potentially biasing its organization and to see if an *in-silico* evolutionary process produces topologies with the functional motif of inhibition between rivalrous elements.

Briefly, we found that not every perturbation necessarily revealed causation. Although data from both lesioning regimes showed similarities, SPA missed a few of the key contributing elements and miss-attributed their causal ranks. Therefore, it provided biased contributions for individual elements, while the MSA captured these nuances more accurately. To further quantify the complex interaction of elements within the system, we used an extension of MSA, here called Pairwise Causal Interaction Analysis (PCIA)[27,28], and found a handful of pairs in which lesioning one unit while the other is perturbed restored the disrupted behavior. Finally, we delved deeper into the inner mechanisms of the network to identify why MSA ranked the units in the given way and what these units do that SPA was insensitive to. We discuss the findings, the limitations of the current approach and outline potential future questions to pursue.

## Results

Our *in-silico* experimental setup was the ATARI arcade game Space Invaders, in which the agent, located at the bottom of the environment, needs to defend itself from aliens descending from the upper part of the screen using laser canons. The main objectives are to stay alive by avoiding alien laser shots and scoring as many points as possible by eliminating aliens. On average, a human subject obtains a score of 1652, and an algorithm that randomly selects actions can reach a score of 148[32]. Other classic algorithms, such as an earlier implementation of a Deep Q-learning Network (DQN), State–Action–Reward–State–Action (SARSA), and a refined DQN, reach 581, 271, and 1976, respectively[32,33].

Instead of training deep networks using backpropagation in a predefined architecture, we evolved a compact network using a Neural Architecture Search (NAS) algorithm called Neuro Evolution of Augmenting Topologies (NEAT)[34]. Briefly, NEAT uses evolutionary principles such as cross-over of genes (network topologies), speciation (preserving novelty), and incremental complexification to find the “fittest” topology. This means the network’s architecture and connectivity are not handcrafted, nor does the algorithm solely optimize connection weights. Instead, the fittest network is evolved with respect to the environmental constraints, in this case, to have the highest score by adjusting its topology according to a set of given limitations, for instance, low probability of adding connections versus higher probability of removing them, see section *Evolutionary optimization*.

In addition to these sets of hyperparameters, to further enforce a compact architecture, we compressed the game frames using a deep auto-encoder and fed our network with two feature vectors (12 features in total, neurons labeled with negative numbers in Fig.1) at each time point. We fed two frames instead of one due to the non-Markovian structure of the game in which only knowing the current position of laser beams does not provide enough information about the beams’ directions.

**Fig.1:**
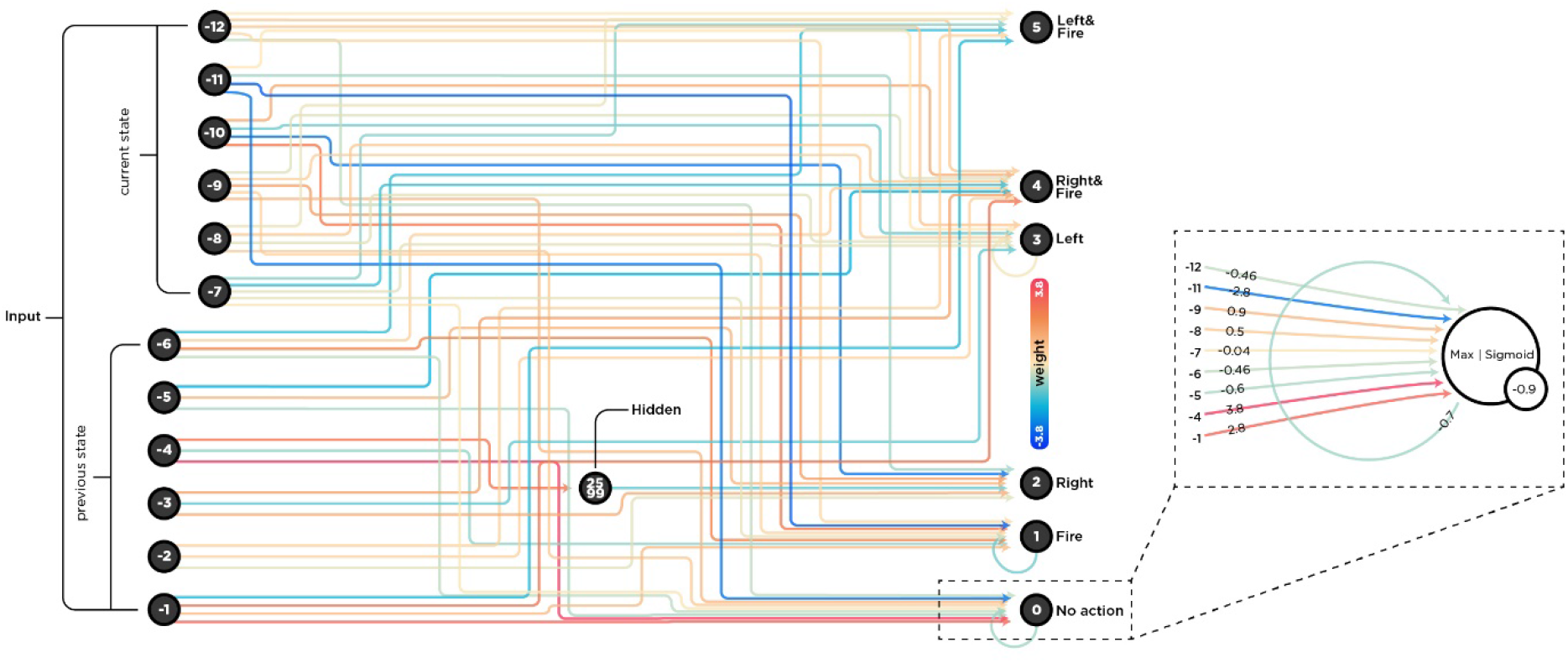
The complete wiring diagram of the evolved ANN. At each time point, the network received a compressed version of the game-state as a vector of 12 features, six features per frame. It then chooses an action from six available actions (output nodes). Due to its importance, which was revealed later in the analysis, we plotted node 0 separately with more information on the right part of the figure. The aggregation function for this node is max, the activation function is a sigmoid function, and the bias is −0.9. Note that these functions are different for each node (see section *Evolutionary optimization*).

On average, our evolved network obtained a score of 337 that is significantly higher than a random agent with a score of 148 (Mann-Whitney U statistics; MWUs = 39542, p-value <0.001, Fig.2). In addition to the random agent and to ensure that the score is not higher merely because of innate topological privileges, we compared the performance with the performance of two control networks. In one, we kept the network as is and made it blind by feeding noise instead of features, and in the other, while the network was receiving game-states, we shuffled the connection weights. Both control networks obtained substantially lower scores, i.e., from 175 (MWUs = 50919, p-value <0.001) to 129 (MWUs = 31157, p-value <0.001), respectively. Altogether, these results show that our compact network did learn the task to some degree and could reach a good enough score (Fig.2) that formed the basis of the subsequent perturbation analyses.

**Fig.2:**
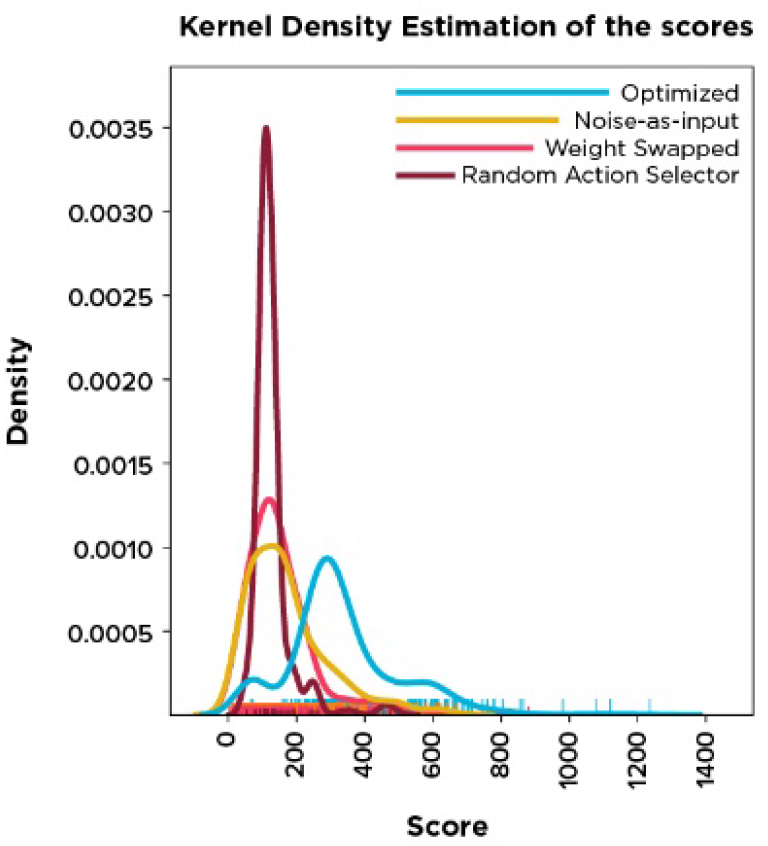
Distribution of performances. Optimized network is the evolved network, which reached a good-enough performance. Noise-as-input is the same network that receives random values drawn from a uniform distribution [0, 1] as input instead of receiving game-states. Weight swapped network receives the game-states while the connection weights are shuffled. Finally, Random action selector is an algorithm that selects a random action, at each timepoint, regardless of the game-states.

### Perturbing all elements, one at a time

After evolving the network, we intervened to see if perturbing elements could reveal their causal importance for the behavior. We first silenced neurons one at a time and ran the simulation with the lesioned network. Conventionally, we searched for neurons, which, when lesioned, resulted in a considerably deteriorated performance, indicating their “necessity” for the behavior. As (Fig.3A) shows, lesioning either of two input neurons −1 and −9 had such a disruptive impact, while individually perturbing most other neurons had a negligible effect on the performance. Interestingly, lesions of two neurons, 4 and −5, improved the performance, suggesting their hindering role during normal functioning.

**Fig.3:**
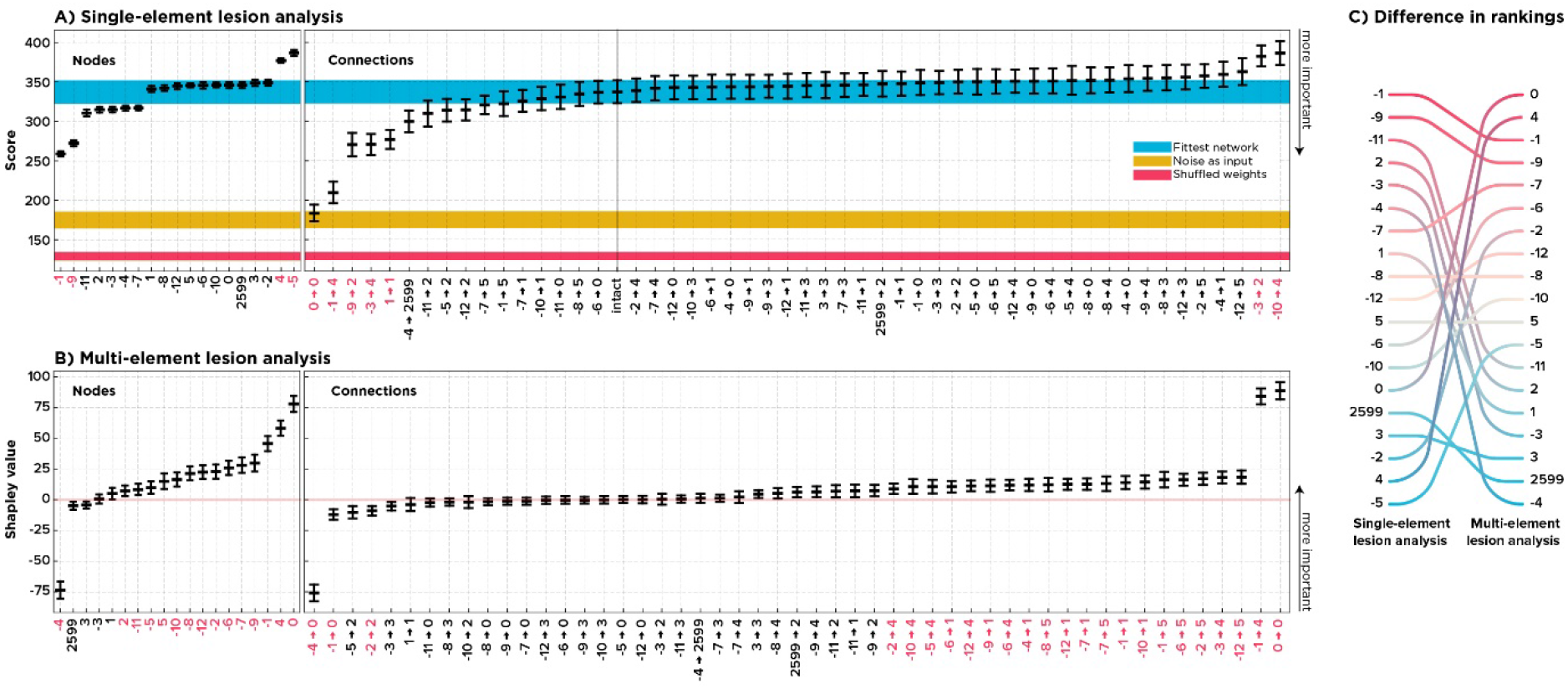
Single-element Perturbation Analysis versus multi-perturbation Shapley value Analysis of the ANN. This figure shows the result and the rank difference derived from a SPA (A; 512 samples per element) versus an MSA (B; 1,000 samples per element). On the left side, the nodes, and in the middle, the connections are sorted according to their inferred average contributions. For SPA, the lowest value means the most influential while the other way around applies to Shapley values, with the highest value means the most critical. Error bars are %95 Confidence Interval (CI; bootstrapped 10,000 times). The blue, yellow, and red strips show the %95 CI of the labeled control networks. Red labels on the x-axis show significant elements (alpha inflation is corrected using Bonferroni correction, see *Statistical inference* in *Materials and methods*). On the right-hand side, the node rankings are compared.

To account for the unique consequences of white matter lesions, also known as disconnection syndromes[11,35], we performed the same lesioning scheme on all connections. We wanted to see if severing individual connections among neurons instead of silencing a whole neuron with all its connections can further localize functional contributions in our ANN. For example, are neurons −1 and −9 essential elements for the behavior of the ANN, or are there connections of these neurons such that the neurons only appear to be critical in the sense that lesioning them perturbed those connections as well? Based on the single-node removal experiment results, we expected to see either no specific connections to be causally crucial, showing that neurons are the actual units of causation or a major disruption in behavior following lesions to the outgoing connections from neurons −1 and −9.

Surprisingly, a loop from neuron 0 to itself (self-loop) appeared to be the most critical element (Fig.3B). This observation indicates that, although SPA of all elements resulted in some degree of coherence by first capturing neurons −1 and −9 as major players and then tracking their importance to connections (−1 → 4) and (−9 → 2), another key aspect is downplayed. If neurons were the essential elements, no single connection lesion would have had such devastating effects, or the critical connections would be associated with the critical neurons. However, lesioning single connections did impact the performance considerably. The critical connection is not a connection from or to the most important neurons but a self-loop of a neuron that itself had a near-zero causal contribution.

To summarize our point, results from the SPA of each neuron indicated that neuron 0 has little impact on the performance while SPA of the self-loop (0 → 0) disrupted the behavior the most. Note that throughout the lesioning experiments, the network was fixed, and its architecture determined its behavior. Therefore, we suspected a more complex interaction among neuron 0’s connections such that lesioning (0 → 0), while those key connections were intact, disrupted the behavior, and lesioning (0 → 0) alongside them had no adverse effect. We suspect those connections to be among other connections of neuron 0 since removing the node virtually perturbed all its connections, which ended in no disruption in the behavior. Put simply, lesioning connection (0 → 0) alone caused the most damage while lesioning neuron 0 with all its 11 connections – including (0 → 0) – did not show any behavioral impairment. In the next section, we describe the MSA algorithm and elaborate on its results.

### Multi-perturbation Shapley value Analysis of all elements

We next adopted a multi-element lesioning approach to perturb all neurons and all connections. We used a rigorous, game-theoretical metric based on the Shapley value (γ) called MSA[27]. To elaborate, the Shapley value accounts for the *“worth”* of an element in the grand coalition of all elements, forming the entire system, in terms of the element’s contribution to the system’s outcome, given its unique added contribution to all possible combinations of coalitions[27–29]. So far, Shapley values are the only values that mathematically proven to satisfy the following axioms[29]:

1. **Symmetry:**If two elements are functionally interchangeable, then their contributions will not differ by their labels.
2. **Null player property:**If an element does not contribute to the given function, its Shapley value is zero.
3. **Additivity:**Summing the contributions of all elements results in the performance of the grand coalition.

As with the SPA framework, this approach aims to find elements that, when lesioned, most strongly impair the behavior. In this case, these elements have the highest Shapley value that is derived from permuting all combinations of multi-site lesions (many elements are lesioned at each time) such that the target element is once included in the lesioned coalition and once excluded from it. In other words, for each permutation, a set of elements are lesioned, the performance is quantified, the target element is then lesioned alongside the other elements in the coalition, and the performance is quantified again. The difference between these two conditions, both negative and positive, is what lesioning an element contributes to that specific group of lesioned elements (Fig.4). Note that the subsets have arbitrarily different sizes, which means the analysis is reduced to SPA if the coalition contains only one element, i.e., the target element, and is expanded to the whole network if the coalition contains all elements. Therefore, while focusing on the importance of one element, MSA incorporates the multivariate influences of lesioning other elements. Averaging over these contributions will then be the Shapley value of the target element, indicating its marginal causal contribution to the system’s performance.

**Fig.4:**
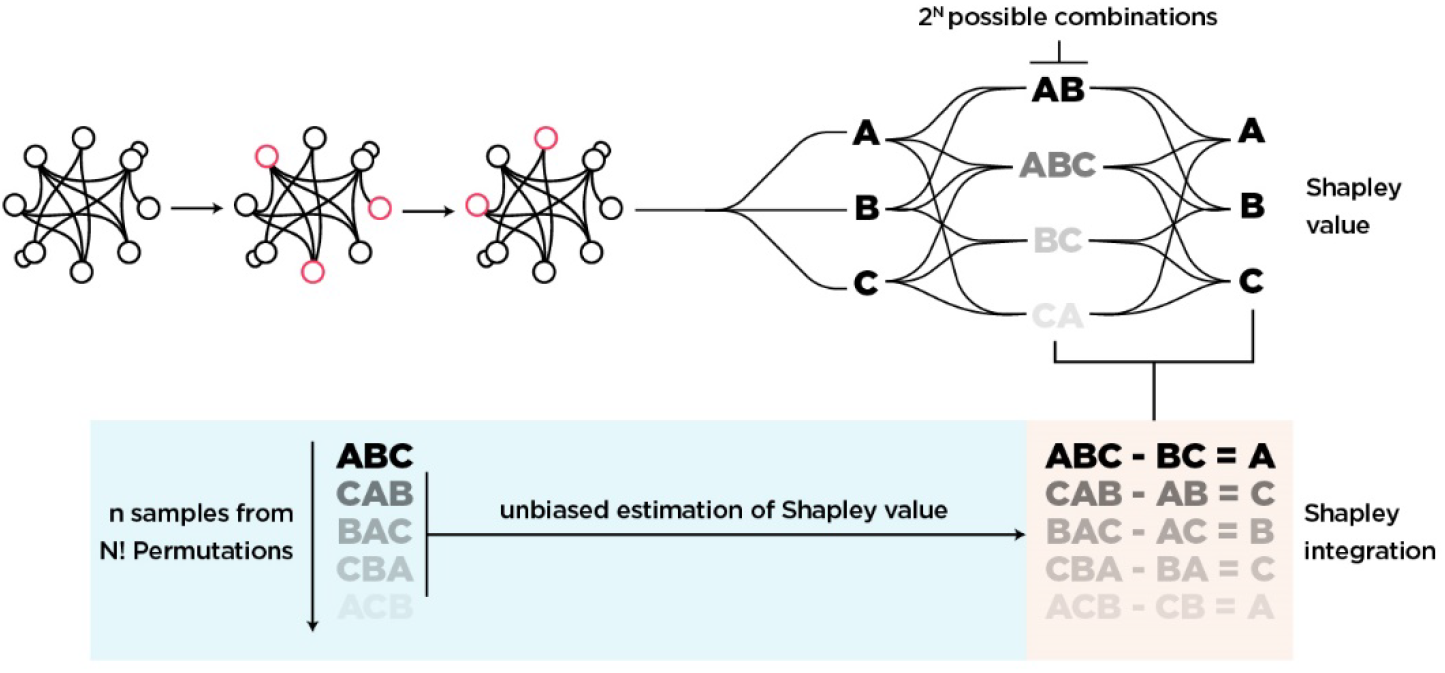
Visual depiction of MSA algorithm. Since there are 2^N^ possible combinations of coalitions, an analytical solution for the Shapley value is computationally prohibitive. Therefore, we sampled 1,000 random permutations from all N! possible orderings and used those to dictate which coalitions to perturb. One sample of Shapley value for any element is then its contribution to one permutation, simply by calculating the score difference of the coalition with the element (e.g., {A, B, C}) and the score of the same coalition without the targeted element (i.e., {A, B} to isolate C). Note that permutations are order-invariant, which means the performance of coalitions {A, B, C} = {C, B, A}.

However, having all possible combinations of subsets explored can be computationally prohibitive in large sets. Therefore, we used an unbiased estimator of the Shapley value that samples coalitions from the space of 2^N^ possible combinations, where N is the number of all elements (see [27] for detailed information).

As mentioned, the Shapley value is additive and thus has an intuitive interpretation in which the highest possible Shapley value is the grand coalition’s worth. For example, for our network, the Shapley value of the overall coalition is 337. This means an element with a Shapley value of 80 accounts for a fraction of 23% of the network’s performance. A negative Shapley value follows the same line of interpretation, that is, an element with a Shapley value of −80 on average prevents the network from an additional 23% increase in performance.

As depicted in (Fig.3B), MSA shows many noncritical nodes and connections, just as the single-site lesion analysis did. Importantly, according to the MSA, neuron 0 is the most influential, followed by many less critical nodes. Interestingly, neuron −4 is assigned a negative Shapley value, indicating its proportionally large and inhibiting contribution to the system. This contradicts the result obtained from SPA that pointed to −5 and 4 to have such an influence (Fig.3C).

As with SPA, we dissected nodes to their connections but this time using MSA to test if we can further track the critical neurons’ causal influence down to their connections. Again, we expected to see either lesioning of no single connection to have drastic effects, indicating a distributed regime of processing in which no lower-level unit is as critical, or to find that there are critical connections, and they correspond to the influential nodes since lesioning a node here is the same as lesioning all its connections.

MSA tracked the importance of −4 to a single connection from −4 to 0, and the same correspondence applies to the elements with the highest Shapley value. The causal contribution of neuron 0, for example, can be attributed to its connection (0 → 0) since besides (0 → 0) and (−4 → 0), other connections of this neuron have negligible contributions (Fig.3B). As a sanity check, we performed the same procedure on the blinded network. Here we expected no element to contribute to the network’s overall performance since, on average, the network had the same baseline score. As shown in (Supplementary Figures 1), this is indeed the case.

The most crucial difference between SPA and MSA was how they ranked connections (0 → 0) and (−4 → 0). Remember, even data from the SPA showed (0 → 0) as the most critical connection. The missing piece was another link to neuron 0 that we suspected to have a Sprague effect-inducing interaction with the self-loop (0 → 0) and the reason was that by perturbing all 11 connections, including (0 → 0), we had no adverse effect. MSA attributed a negative Shapley value to the connection (−4 → 0), while SPA assigned minor importance to this connection. This discrepancy aligns with the Sprague effect’s essence since at least two elements are required to be lesioned for such a phenomenon to emerge.

Altogether, MSA and SPA found key elements to be a small and localized set. MSA dissociated these and assigned the negative contribution to neuron −4 while SPA missed it. While SPA excluded neuron 0, MSA ranked it as the most critical neuron and further dissected this importance to the self-loop. It then showed that the incoming connection from −4 is the possible answer to why lesioning neuron 0 has a near-zero impact.

### Impact of lesioning on functional connectivity

In addition to their direct impact on the behavior of a system, lesions may also disrupt functional connectivity (FC), and different features of the impact on FC are associated with behavioral performance. Thus, FC forms a bridge, or ‘intermediate phenotype’ from structure to function and behavior [35–38]. It was shown that lesions of critical brain regions in terms of FC, such as hubs, have a greater impact on the dynamics of the whole brain[37]. To explore this aspect in our *in-silico* model, we first calculated the FC of the intact network using Pearson’s correlation. We then employed a SPA framework for all units, that is, nodes and connections. To quantify the impact of lesioning individual elements on global FC, we calculated the element-wise differences between intact and lesioned FC matrices. The absolute sum of the resulted difference matrix was considered as the Impact of lesioning on Functional Connectivity (IFC; Fig.5). A larger IFC results from a greater difference between FC of the intact network and FC of the lesioned network and intuitively indicates the importance of elements, this time by their contribution to overall functional connectivity instead of performance.

**Fig.5:**
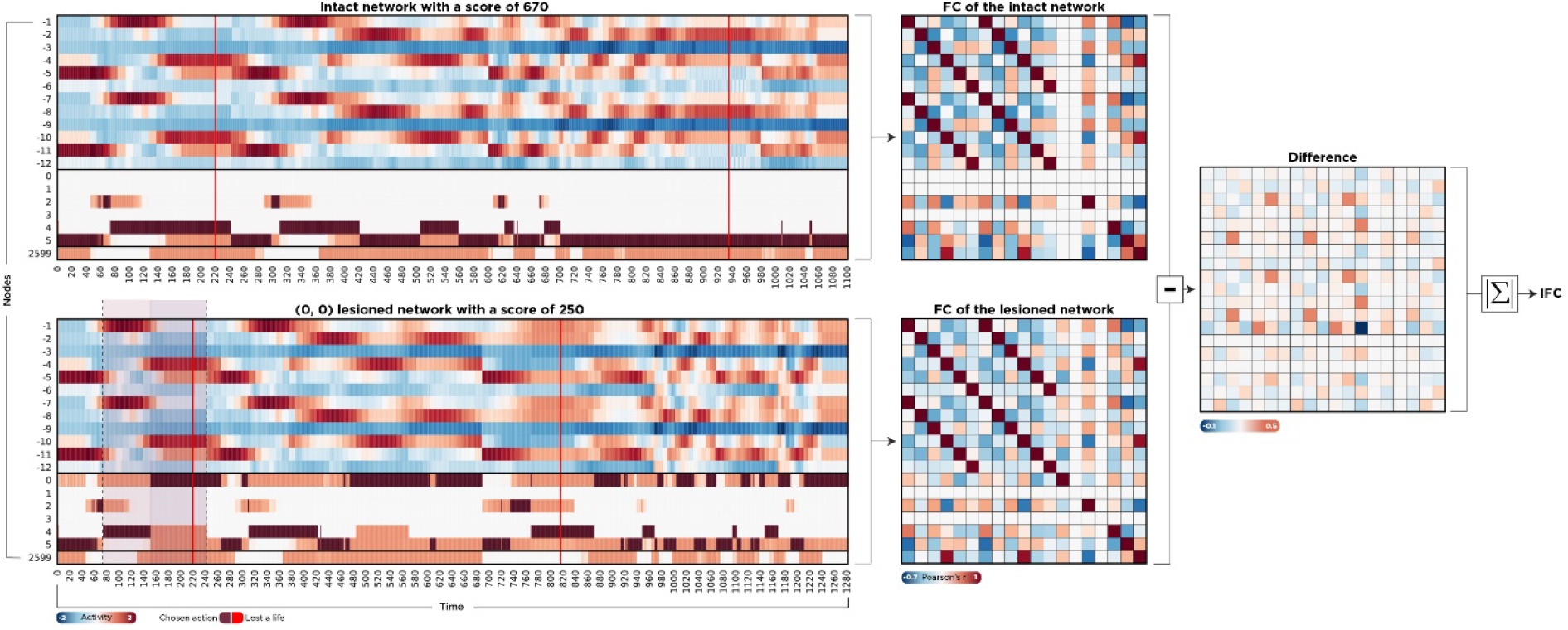
Calculating the impact of lesions on functional connectivity. We recorded the activity of all neurons to compute the functional connectivity of the network. We exhaustively perturbed all units one by one and compared the element-wise differences between intact and lesioned FC matrices. The absolute sum of this difference matrix (IFC) quantifies how much a lesion caused the network dynamics to deviate from its uninterrupted state. On the left-hand side, the activity of two scenarios is depicted. In the upper timeline, the network is intact, and the score is 670, while in the lower timeline, the feedback loop (0 → 0) is lesioned, leading to a drastic decrease in performance. Red vertical lines showed when the agent was shot and lost a life. Brown cells indicate the chosen action, and the dashed window is the same time window that we zoomed in further in the section *Understanding the Paradoxical lesion.*

Interestingly, IFC is negatively correlated with both nodal and connection perturbation scenarios, corroborating previous findings (Fig.6). However, IFC is not associated with Shapley values of these elements. This means that, although SPA has internal coherency by identifying units that, perturbed one by one, have the largest effect on both functional connectivity and the agent’s performance, these units are not the same as those captured by an MSA framework. In other words, the bridge is formed. However, as shown in Fig.3, the actual players remained obscure. We show why the rankings differ and propose a possible underlying mechanism that accounts for this discrepancy in the next two sections.

**Fig.6:**
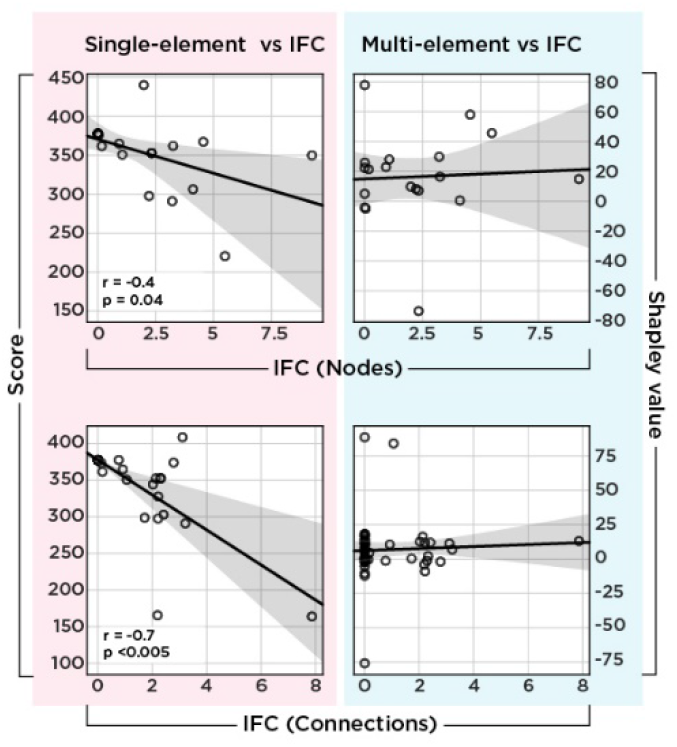
Correlation between IFC and single-site lesioning scheme. The upper left scatterplot shows the relationship between the impact of the SPA of nodes on functional connectivity and the agent’s performance. The lower left scatterplots show the same relationship but for each connection. Both show a negative correlation, which means the larger the impact on functional connectivity, the lower the performance. However, this relationship is absent from the right-hand side that compares the Shapley value of each element with their IFC. As with the left-hand side, the x-axis shows the IFC of nodes (upper plot) and connections (lower plots), while here, the y-axis represents Shapley value instead of raw performance.

### Quantifying complex interactions between causal building blocks

In previous sections, we presented two causal rankings of elements from the same ground-truth neural network model, one using a SPA framework and the other using MSA (Fig3.C). We found that the changes in the inner dynamics of the system perturbed using SPA support this approach’s ranking, which mistakenly adds more certainty to the accuracy of the approach in finding critical units. Here we show why these rankings differ by measuring the complex interactions of units. Although MSA is a multivariate approach that accounts for a large variety of combinations of units, it eventually describes the system in terms of how much, averaged over all combinations with other units, *single units* contribute to the output. In other words, it isolates the average *individual* contributions and not the nature of their interactions. Using an extension of MSA, here called PCIA, we formalized and then quantified these interactions since the causal influence of one element is intertwined with the state of others.

At its core, PCIA is a chain of multiple MSAs in different conditions. To elaborate, quantifying the complex pairwise interaction of two elements *i* and *j* requires first to calculate the Shapley value of them both as a single compound element *γ*_(*ij*)_, followed by the Shapley value of each one given the other is perturbed 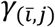 and 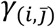 respectively. As Fig.7 shows, subtracting all three provides an interaction term that, if positive, indicates *“synergy”* between the pair and, if negative, shows *“redundancy”* or functional overlap. In other words, PCIA quantifies how much the causal contribution of a pair of units is bigger or smaller than the sum of their individual contributions.

**Fig.7:**
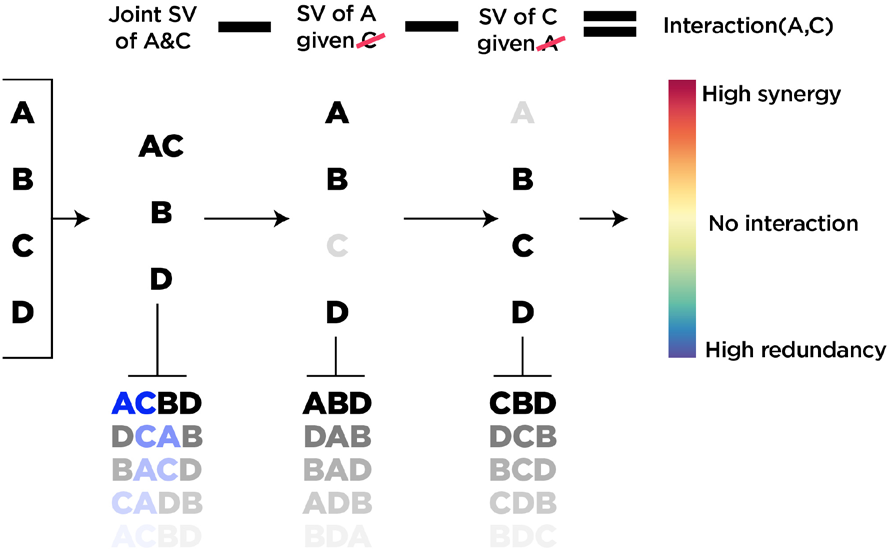
Visual depiction of the PCIA algorithm. At its core, PCIA comprises multiple MSAs. We first start with calculating the joint contribution of two elements, followed by the contribution of each, given the other is perturbed. The interaction term is then calculated by subtracting these values from each other, indicating how much the joint contribution of a pair of elements is bigger or smaller than the sum of their individual contributions. Like MSA, permutations are order-invariant.

Since PCIA involves the calculation of multiple MSAs, it is computationally even more expensive. Therefore, we focused on the connections, and to calculate all pairs of them, we sampled 100 permutations per element instead of 1000, as in the case of MSA.

The results are shown in Fig.8, and as quickly stands out, there is a strong synergy between two elements (0 → 0) and (−4 → 0), followed by a handful of strongly redundant and many minuscule interactions in both directions. Therefore, the results from this analysis provide more evidence for (0 → 0) and (−4 → 0) to have a unique form of interaction, which we next investigate with respect to whether it is a paradoxical lesion-effect.

**Fig.8:**
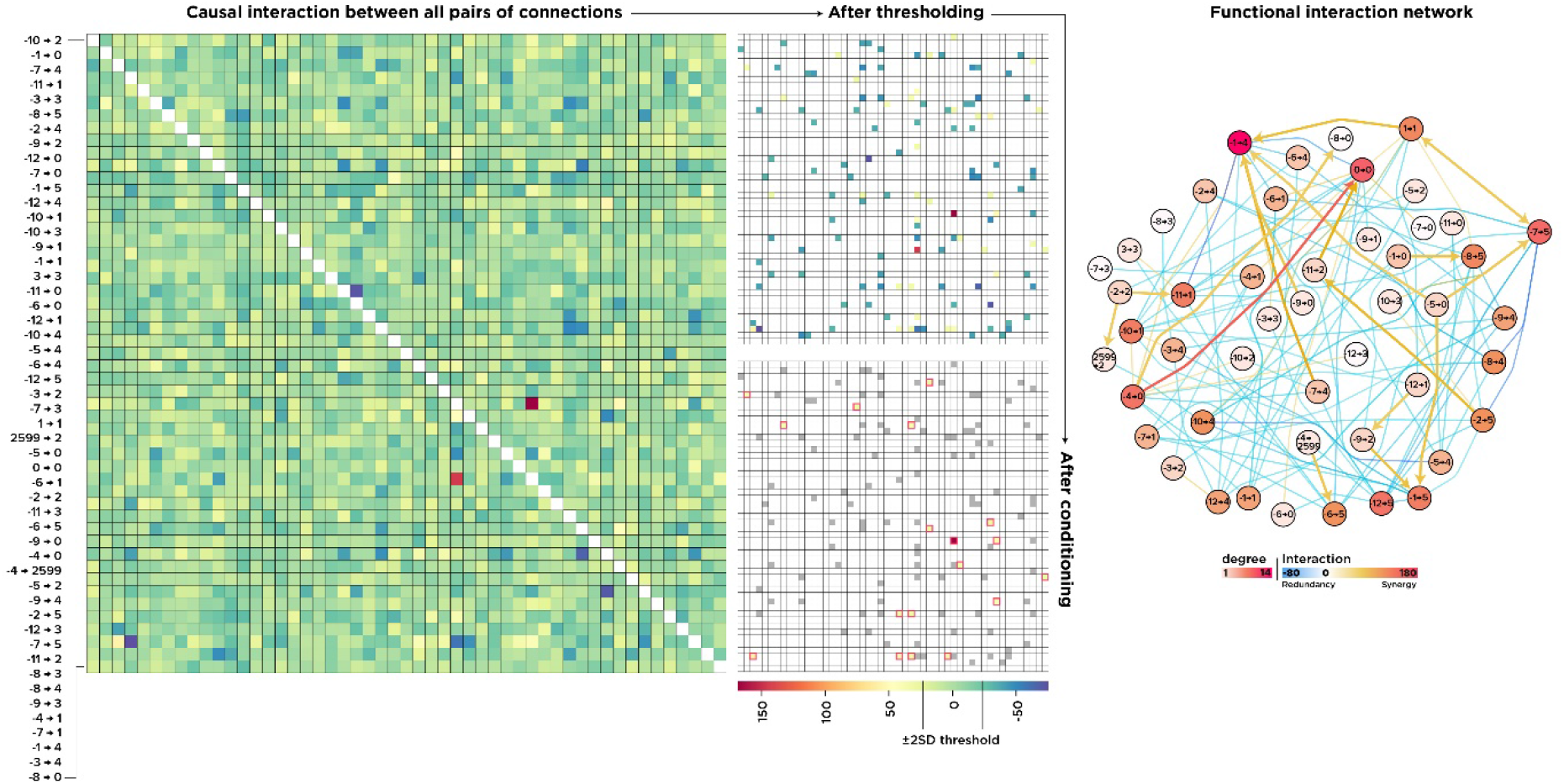
Pairwise interactions among all connections. An interaction matrix resulted from the PCIA procedure in which warmer colors show greater synergy and cooler colors indicate functional overlap (left). We then excluded ±2 SD and applied the “Sprague effect” condition to the thresholded matrix (middle). On the right-hand side, we plotted the interaction network in which the nodes represent connections in the actual network, and the edges are interactions among them. Arrows show paradoxical-lesion effects ( i → j ).

To do so, we formalized the Sprague effect as the difference between the average importance of element *i* given the state of the element *j*. Specifically, the Sprague effect is defined as a scenario in which element *i* has a negative Shapley value when element *j* is perturbed 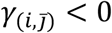, thus hindering the performance and has a positive contribution when *j* is intact 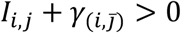. Put simply, on average, element *i* disrupts the performance if element *j* is intact and improves if *j* is lesioned[28,31].

To reduce the number of false-positive findings, we looked for this condition among a smaller set of pairs with an interaction term above and below two standard deviations of the mean. The results are shown in Fig.8, with connections indicating the interactions and arrows depicting a Sprague effect between two elements (the stem of the arrow indicates the element *i* that has a negative contribution when the pointed element *j* is lesioned.) As depicted, we found many paradoxical lesion effects predominantly among synergistic interactions, with the interaction between (0 → 0) and (−4 → 0) being the most prominent one. This network is a higher order “functional/interaction network” in which its nodes represent connections in the “structural/actual network”.

To summarize this part, we first quantified how much two elements’ causal importance is larger or smaller than the sum of the individual elements. We then used this metric to classify the modulatory effect of each element on the others, with a focus on paradoxical modulations, and found a handful of elements in which lesioning one, while the other is perturbed, restored the performance. The connections (0 → 0) and (−4 → 0) had the highest synergy, meaning that, together as a whole, they functionally contribute much more than their summed individual contributions. This unique synergy is also a paradoxical-lesion effect in that lesioning (0 → 0) alone disrupts the performance while lesioning it alongside (−4 → 0) restores it. Note that the metric captures what a SPA framework is insensitive to, specifically, complex pairwise causal interactions. In other words, PCIA is built upon MSA that, as seen, extends SPA to lesioning combinations of elements, and here, it is systematically bundled to quantify complex multivariate relationships that elements might have. These interactions and insensitivity of SPA to them are what, we believe, eventually leads to misattributing key elements in their ranking of causally critical units. By focusing on the two connections (0 → 0) and (−4 → 0) in the next section, we show paradoxical lesion effects might not be that unlikely and, quite contrary, they might be a direct result of perturbing a simple and ubiquitous motif of connectivity, which explains why we found many such paradoxical effects in this analysis.

### Understanding the Paradoxical Lesion

The Sprague effect was first discovered in cats and later in humans, with its underlying mechanisms still partly elusive[19,20]. One current theory suggests the phenomenon is caused by a reduction of inhibition from a functionally competing region, and the deficit reverses when both are lesioned[22]. To see if this is the case in our network, we focused on the two most prominent units (0 → 0) and (−4 → 0). Note that the SPA also ranked (0 → 0) among the most critical connections. However, (−4 → 0) was only captured by MSA and was the only unit with a large negative Shapley value. The top plot in Fig.9 shows the activity of two input units, −1 and −4, over the trial in which (0 → 0) is lesioned (also see Fig.5). A Pearson’s correlation analysis shows they are negatively correlated.

**Fig.9:**
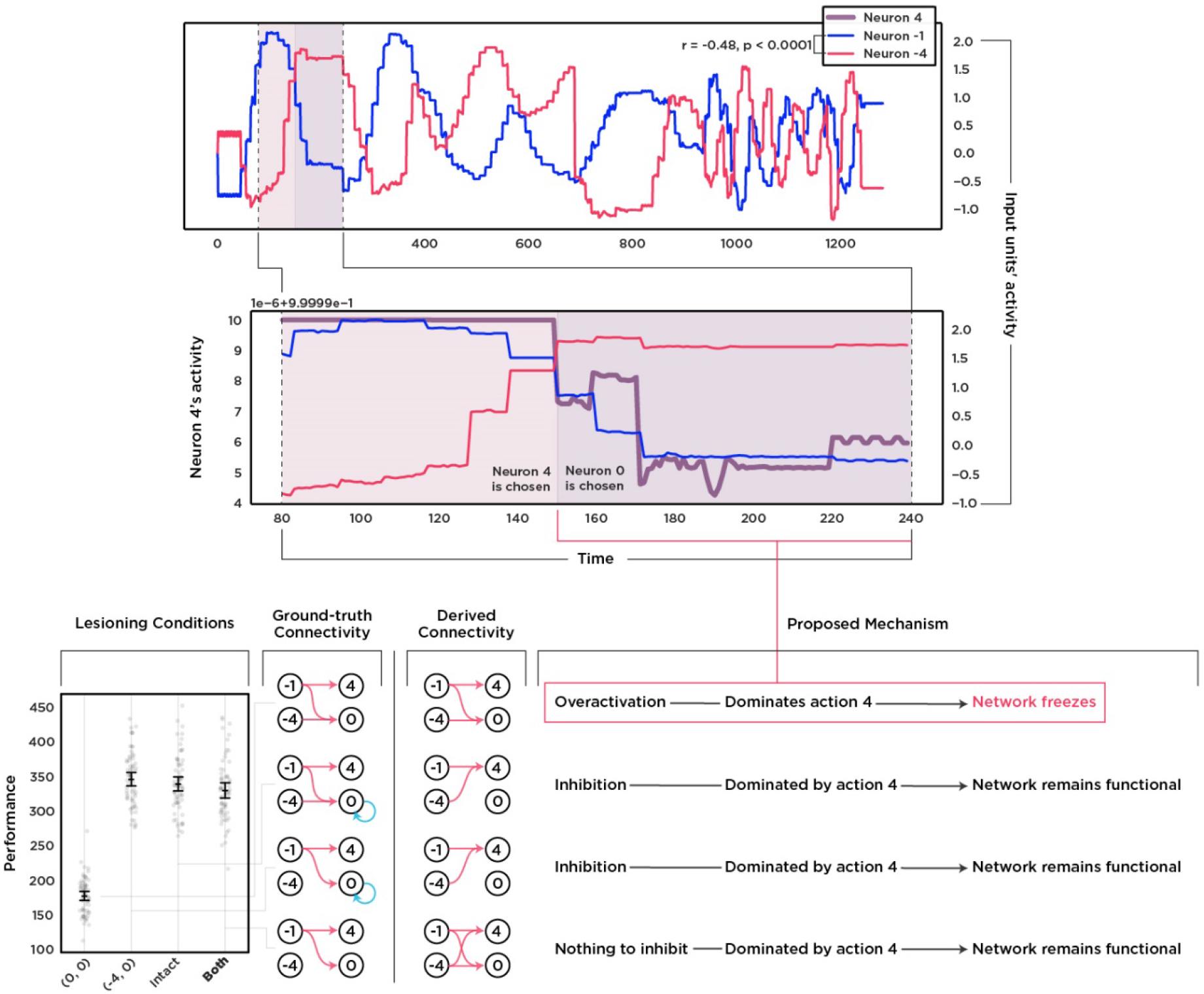
Focusing on the critical elements discovered by MSA. The upper timeline shows the negative correlation between the activity of two input units −1 and −4. This anticorrelation leads to competition between downstream units 4 and 0. In an intact network, unit 4 is dominant due to the inhibitory feedback loop of unit 0. The middle plot shows how unit 4 loses to 0 after the inhibitory loop is lesioned since it is tightly following the input from −1 while neuron 0 is driven by the input from −4. The bottom-left part shows the implications of this rivalry on the performance and how it produces the paradoxical lesion effect. Lesioning the feedback loop disrupts the performance while lesioning it alongside the input from −4 restores the deficit since neuron 0 stays dominated. The bottom-middle part shows the discrepancy between the actual flow of information and the inferred flow by an mTE analysis. Notice the absence of connection between −4 to 0 in the intact network due to the self-inhibition of the target neuron.

Unit −1 is one of the key input units to neuron 4, which itself is one of the most frequently chosen actions by the intact network. Input unit −4, however, has a major influence on neuron 0 (Fig.1) that is inhibited by the negative feedback loop, causing neuron 0 to be silent in the intact network. Since neuron 0 is the action “no action,” the intact network always chooses an action, either 4 (right and fire) or 5 (left and fire). As depicted in Fig.5, lesioning the feedback loop disrupts the inhibition that leads to hyperactivation of neuron 0. Interestingly, although neuron 0 is now competing with neuron 4, it takes roughly 150 timesteps to be selected as the chosen action. The middle part of (Fig9) shows how the decaying activity of unit −1 at around that timepoint causes neuron 4’s activity to follow and eventually lose to neuron 0 in the lesioned network. Naturally, the behavioral consequence of excessively choosing “no action” is gaining a substantially lower score. By lesioning the input from −4 to 0 with or without the feedback loop, the node never reaches the critical threshold to dominate other actions, and thus, in both conditions, the performance remains uninterrupted (Fig.9).

Altogether by looking deeper into the inner dynamics of these units that MSA distinguished, we see a simple motif of connectivity among only four units is enough to produce a paradoxical lesion effect. The key nodes are neurons −4 and 0; the key connections are (0 → 0) and (−4 → 0). The input from −4 to unit 0 has a large negative Shapley value because in coalitions without (0 → 0), it over-activates neuron 0 and causes the network to freeze. The feedback loop (0 → 0) has a high positive Shapley value because it prevents this over-activation, and removing it causes the network to freeze. Interestingly, the input from −1 to 4 has the next highest Shapley value because, without it, unit 4 is dominated by other units, especially an over-activated “no action” node.

A crucial side effect of the functional contribution of silenced nodes is that it becomes very difficult to infer their causal relationship relying on time-series analyses. Here we used a Multivariate Transfer Entropy (mTE) analysis on the four key players in three lesioning conditions and the intact network to see how well they infer information flow in the circuit. As Fig.9 shows, in conditions that neuron 0 is inhibited, mTE missed the information flow even though the node receives input from both −1 and −4.

To conclude this section, we showed that a paradoxical lesion effect could emerge from a simple inhibitory motif. In our case, the inhibition is a negative feedback loop, and the competition is between two output neurons, 4 and 0. We then used mTE analysis to infer the causal relationships that resulted in a critical relationship between −4 and 0 to be overlooked. This shows the necessity of employing systematic lesioning alongside methods relying, for example, on the analysis of time-series dynamics. Altogether, we show that, even in a simple agent, finding which elements are causally relevant for behavior and how, is extremely difficult to answer with confidence. In the next section, we discuss our results, limitations, and future improvements.

## Discussion

In this work, we defined causation not as events prior to effects nor as entities that raise their probability of occurrence but as *contributors to the effect*. Having this definition of causation, we aimed to understand an ANN in terms of its components’ causal influence over its performance. We initially lesioned both its neurons and connections one at a time. We then showed that even with such an exhaustive analysis, which is yet to be reached *in-vivo*, the results are persistently biased. We then formed a bridge from structure to function and eventually to behavior by measuring the impact of single-element lesioning on global functional connectivity. The results supported the ranking from the SPA and added more confidence to the biased conclusion about which units are critical. In other words, our SPA confirms the results from Jonas and Kording’s work[26], and we, too, ended up with structured but biased results.

We then used MSA, a rigorous game-theoretical algorithm, and found the causal ranking to be different. For example, neuron 0 had the highest causal contribution even though it has no major role according to SPA. MSA then identified crucial connections and ranked (0 → 0) the most causally important. It also found neuron −4 to hinder the system and tracked the disruptive element to be the connection (−4 → 0). Next, using an extension of MSA, we first quantified the complex pairwise interaction of all causal building blocks (connections) and, after formalizing the Sprague effect, found lesioning connections (0 → 0) and (−4 → 0) to have such an effect. Lastly, we looked into these two units and found the rivalrous interaction to be the potential mechanism.

Two points to bear in mind are 1. our network was fixed throughout the experiments, leaving no space for plasticity, and 2. the network is a simple ANN with no excitatory-to-inhibitory synaptic dynamics. It is indeed possible that these physiological mechanisms underlie paradoxical lesion effects in the living brain[20]. However, we did not include them in our model; therefore, we believe the paradoxical effects observed here result from none of these mechanisms. We found functional inhibition between competing units sufficient to produce a Sprague effect, as also investigated before ([22,23] and see [31] for a fixed artificial network).

Besides that, further research is needed to compare different mechanisms using biologically plausible neural network models since understanding the phenomenon also relies on different analytical approaches as we used PCIA while Sajid et al. [25], for example, used a dual-lesion scheme. On the same line, since our results point towards a type of interaction that is possibly rooted in the pattern of connectivity in a very rudimentary system compared to the human brain, comparative studies can shine a light on how deep the motif is embedded in the evolution of nervous systems and what, if there is any, are the adaptive values. Interestingly, in our model, the motif is more costly and sub-optimal because instead of simply removing the input from −4 to 0, the evolutionary process added a negative feedback loop to cancel the disruptive influence producing the motif that leads to a paradoxical lesion effect.

Overall, our results, first and foremost, show the inferential limitations of SPA. We believe many aspects of a system can indeed be investigated and understood by lesioning its elements one at a time. However, it is important to know which aspects cannot. The example we used was the Sprague effect, which was argued to be either noise or exceptional[39]. We speculate that if our compact network with 19 neurons and 51 connections evolved at least one of such effects, then it might not be a rare event (as also argued in [5]) but an indication of complex multivariate functional motifs of computation as proposed in [24].

A substantial challenge in depicting a mechanistic blueprint of any system is to have a solid causal understanding of it. The conventional approach perturbs its elements and pinpoints those resulting in a disrupted behavior[10]. These elements were then called necessary causes of the observed effect since they serve as critical substrates for an intact behavior[40]. However, there have been arguments against the classification of neural components as such (see [41–43]). Supporting those arguments, we propose that one step towards a solid causal understanding of the brain is to instead *quantify the degree* to which its neural elements contribute to cognition and behavior.

To put it into perspective, for behaviors to emerge, many neural circuits coordinate, cooperate, and form coalitions that boiled down to a single “necessary” entity, resulting in losing crucial information of the brain’s inner workings[6,42]. This was the case in the contributions we derived from SPA. Thus, we used MSA to capture the whole spectrum of causation instead. Shapley value results from a mathematically sound analysis of all possible combinations in which units can form coalitions and produce the behavior, either flawlessly or disrupted. In its essence, Shapley value is the *fair* share of the elements in producing the function so that the most important elements assigned the highest share followed by a continuum of importance to zero for independent elements and negative values for hindering ones. Therefore, it provides a rigorous and intuitive way that neural elements can be ranked according to their causal contributions to the under-investigated behavior.

Although powerful and intuitive, it is important to emphasize what Shapley value is not (see [44] for a more technical perspective). For example, Shapley value by default does not reveal mechanisms neither it shows what computations were done by individual elements. It shows *how much each element is functionally contributing to the underlying mechanistic processes.* As mentioned, we believe this is *the first step* towards a more comprehensive mechanistic description of the brain, illuminating which elements to focus on next. We, too, did so by focusing on the few key elements that summarize why the intact and lesioned networks behave such and why MSA chooses these units as causally relevant.

We can gain profound insight into the system by incorporating MSA in more complicated analytical pipelines. For instance, the elements’ Shapley value will vary according to the behavior of interest. In principle, one can produce a multidimensional map of each element, knowing how much they are involved in various behaviors. This is relevant for neuroscience since it is shown that brain regions are multifunctional and play different roles in different coalitions[45,46]. Having negligible Shapley value in a task is not an indication of inutility but an indicator of independence since the same element might have a considerably large share in the emergence of a different function. This feature can be used to decompose and dissociate roles that neural elements play in different tasks. For example, in this study, we used the system’s overall performance as our metric that is the product of many behavioral primitives and found specific elements to be the most critical.

Further analyses can decompose the behavior, as we did with the system itself from its nodes to its connections, and calculate the causal share of the elements in each behavioral component, such as, in our case, actions that construct the learned strategy. Therefore, we can expand our knowledge of how elements dynamically form coalitions to solve sub-tasks of the given task, providing a detailed description of the system’s inner mechanism. In other words, given an elegant experiment in which behavior and its components can be measured, Shapley value is a robust method to unravel how neuronal units adaptively join communities and produce hierarchies in the brain. We believe MSA is a powerful tool that can be used to understand the system far deeper than we attempted to do here since, as described above, it has many favorable features and provides intuitive results.

In this work, we used a version of Evolutionary Autonomous Agent models advocated by [47] to be nifty tools for neuroscientists. Using NEAT, we allowed the network’s topology to evolve with respect to the environmental constraints instead of modeling the architecture ourselves and optimize the weights or readout units. This way, we liberated ourselves from further assumptions about the network’s connectivity and structure. It is important to note that NEAT itself produces simple networks that can do simple things. However, more advanced NAS algorithms such as Hyper-NEAT[48] are gaining popularity in the AI community since they produce larger networks that are not limited to the experimenter’s design[49].

Interestingly, in some cases, genetic algorithms rival the conventional Gradient Descent-based methods in non-trivial tasks[50]. This shows a potential role for such algorithms in neuroscience since one can evolve arbitrary architectures to solve an ecologically valid task, e.g., foraging in a patchy environment[51], and compare their topological features with brains evolved in such environments. This extends the toolboxes available to computational neuroscientists, neuroethologists, and behavioral ecologists to more realistic *in-silico* models and experiments.

More cognitively and clinically oriented, *in-silico* multi-element lesioning experiments can be used as a predictive tool to guide non-invasive brain stimulation experiments. For example, human brain connectivity can be used as the backbones of ANNs trained to solve cognitive tasks[52–55]. These connectivity-aware ANNs can then be investigated thoroughly using MSA to predict the critical regions and the corresponding behavioral deficits. The predictions further can be used as testable hypotheses about which regions to perturb *in-vivo*. In other words, connectivity-aware ANNs, neural network models of cognitive processes[56], and large-scale models of functioning brains[57] can add a unique value to the repertoire of ground-truth models to test brain-mapping tools and their limitations (Fig.10).

**Fig.10:**
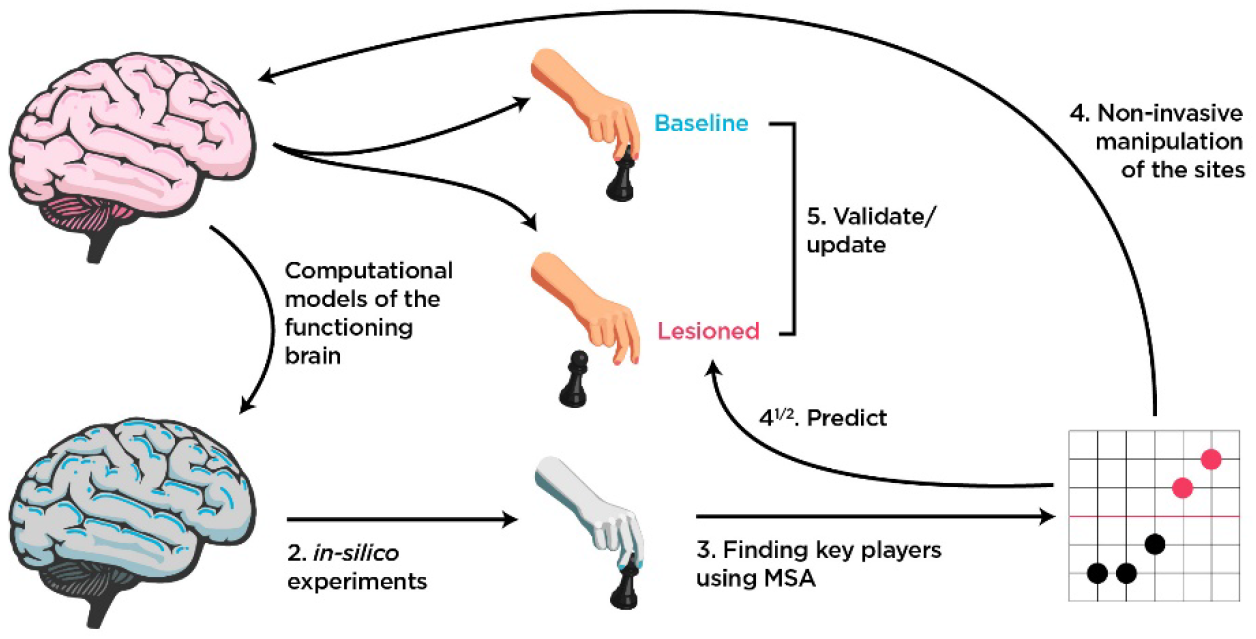
How MSA can be incorporated into the causal brain-mapping toolbox. Since multiple *in-vivo* lesioning is beyond the reach, we suggest connectivity-aware or neural network models of functioning brains to fill the gap. *In-silico* experiments then can be performed to predict both key elements and their contributions to the behavior. These predictions can then be tested *in-vivo* by the method of choice.

The main limitation that is needed to be addressed is MSA’s computational complexity. Having an analytical solution for Shapley values of large systems is an NP-complete problem[58]. Therefore, heuristics[59], predictors[30], and estimators[28,60] are used and are under development to solve this issue. Interestingly, Shapley value has found a unique spot in the field of explainable machine learning[61] and is used to understand deeper and more complicated neural network architectures[61], prune the unnecessary elements[62], and even correct biased networks[60].

Another limitation here is thresholding the “interaction matrix” (Fig.8). As mentioned, even reliably estimating all elements’ pairwise interaction can quickly become impossible since the number of elements is now squared, and three Shapley values are needed for each interaction. Therefore, we reduced the number of samples from 1000 to 100, which means less certainty in the estimated results. To partially account for this problem, we excluded two standard deviations above and below the mean. A decision that directly influences the number of discovered paradoxical-lesion effects. Therefore, a central interest is to address this issue using either better thresholding criteria or estimation methods.

## Conclusion

A common way of characterizing the causal contributions of elements in a system is to perturb them and measure the effect. We showed that not every perturbation reveals causation since lesioning elements, one at a time, produced coherent but biased results. We then used MSA and captured the crucial details missed when we lesioned each site independently. We then found a motif of functional inhibition among competing units to be the underlying mechanism of the paradoxical lesion effects in our network. We believe this effect is the main contributor to the bias in a single-site lesion analysis since, by definition, it emerges from a condition with at least two lesions. This showed that even compact ANNs show surprising complexity that is needed to be addressed to have a step towards a comprehensive causal picture of the system.

Lastly, in the context of rapidly evolving sophisticated uni-and-multivariate brain-mapping methods, we advocate using *in-silico* experiments, and ground-truth models, especially neural network models verify fundamental assumptions, technical limitations, and extent of interpretations of these methods.

## Materials and methods

In this section, we explain the methods and materials used in this research. The codes and generated datasets are publicly available in the following repository:

https://github.com/kuffmode/ANNLesionAnalysis

Briefly, we first trained a deep autoencoder to compress the screen pixels to a handful of features per frame. We then evolved a controller network to, based on these features, choose a proper action. After having both networks, we started the lesioning experiments.

### Evolutionary optimization

We used the NEAT-Python toolbox[63] to evolve a network from an initial stage of randomly connected 12 input and six output nodes. During the evolutionary process, the algorithm was optimizing many parameters, including the choice of activation functions, aggregation functions, adding or removing hidden neurons, adjusting connection weights and node biases, and adding or removing connections (see Table1 for a summary and the file `AEconfig-SI.txt` for the complete list of hyperparameters). There were no restrictions on the connectivity pattern so that a recurrent architecture could evolve from the initial feed-forward stage. We chose the probability of removing connections to be slightly higher than adding (0.6 versus 0.5) to encourage sparsity. We then ran the evolutionary processes 32 times to have 32 candidates. Each time the process ended either after 128 trials or one member reached the fitness criterion of 1200 points. In each trial, the generation comprised of 128 members that were instantiated from the same initial stage and would play the ATARI game independently. After each step, the algorithm mutated the genome according to the given probabilities and performed the cross-over among the top %30 networks to produce the next 128 members. At the end of the training phase, 32 candidate networks reached either the generation limit or the fitness criterion. We then chose the one with the highest score of 1300 points to move forward with the lesion experiments.

**Table1:**
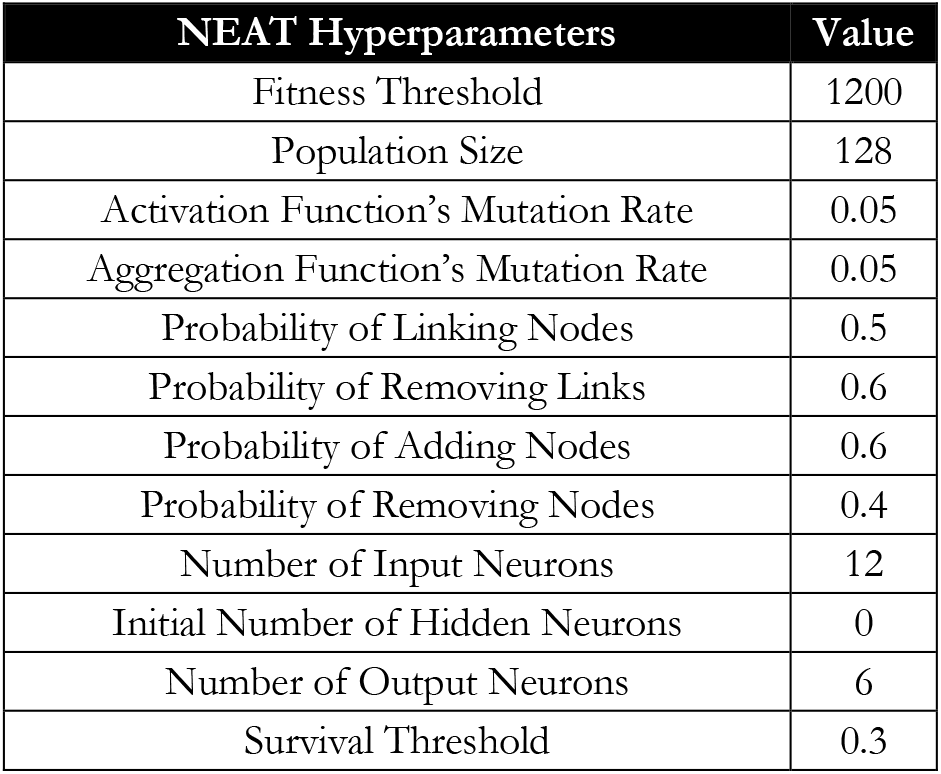
A summary of relevant NEAT hyperparameters. NEAT produces a large variety of networks, all from a set of constraints and probabilities. Since our goal was to produce a good-enough network, we did not tune these parameters for maximum performance and either used the default values or adjusted them according to the experimental objectives, e.g., sparse connectivity.

| NEAT Hyperparameters | Value |
| --- | --- |
| Fitness Threshold | 1200 |
| Population Size | 128 |
| Activation Function's Mutation Rate | 0.05 |
| Aggregation Function's Mutation Rate | 0.05 |
| Probability of Linking Nodes | 0.5 |
| Probability of Removing Links | 0.6 |
| Probability of Adding Nodes | 0.6 |
| Probability of Removing Nodes | 0.4 |
| Number of Input Neurons | 12 |
| Initial Number of Hidden Neurons | 0 |
| Number of Output Neurons | 6 |
| Survival Threshold | 0.3 |

### The preprocessing steps and the Autoencoder

We used OpenAI Gym[64] ATARI environment as our game environment. The game screen generates an array with the size of (210, 160, 3); since the screen is 210 × 160 pixels, each contains three color values, red, green, and blue. Throughout the whole work, the pixels passed through a preprocessing pipeline first that would:

1. crop-out the unrelated parts of the screen such as scores and the ground,
2. convert colors to the monochrome gray-scale, therefore reducing the 3D space of red, green, and blue values to one intensity value representing brightness of each pixel,
3. binarize the pixel values to either an *“on pixel”* or *“off pixel”*,
4. and finally, flatten the outcome into a vector with a size of 2679 pixels.

This vector represented the game with a series of zeros and ones that were then fed to the Autoencoder (Fig.11). The Autoencoder was a Keras model[65] trained independently from the controller network. We first recorded 43,200 frames from the game played by a random agent, shuffled the frame orders, and used 28,800 frames (≈%65 of the dataset) to train and the rest for testing the Autoencoder. The architecture was designed with four encoding layers and four decoding, and a bottleneck of six features.

**Fig.11:**
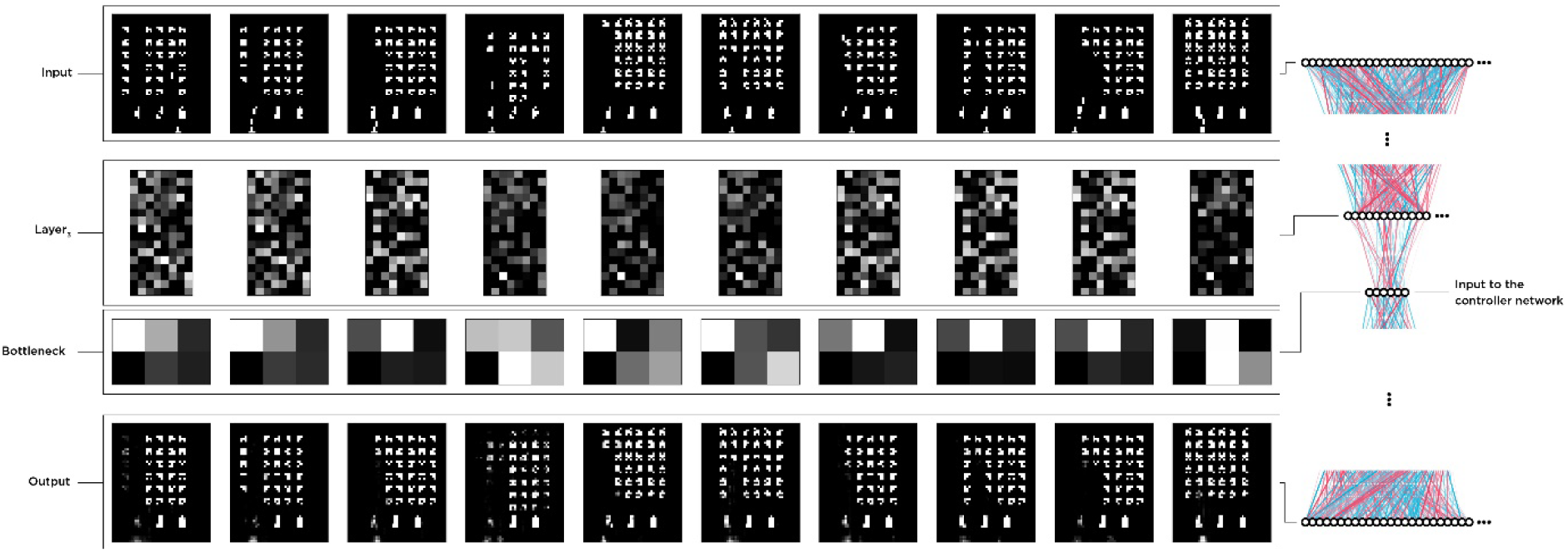
Visualization of the Autoencoder’s inputs, latent features, and decoded outputs. The Autoencoder was trained separately from the controller and received recorded frames from a random action selector agent. We then used the encoder half to reduce the pixel space to six features per frame and fed the controller with two feature vectors.

We used the ADAM optimizer, a binary cross-entropy loss function, 64 epochs, and a batch size of 512. Since input frames are binarized, we used Rectified Linear Unit (ReLU) activation functions for all layers except the last decoding layer, for which we used a Sigmoid function instead. After the training session and accuracy of ≈%98.8, we kept the encoder network and fed the latent space to the controller network throughout all experiments and the evolutionary process of the controller (Fig.11). Together the Autoencoder and the controller network formed our agent. However, we did not perturb the Autoencoder and focused solely on the controller during the experiments.

### Lesion Analysis

We first pruned our network by pruning the already “disabled” connections. Briefly, connections in the network are either enable, meaning they multiply the incoming value with the weights and pass it to the receiver node, or disabled that pass zero. During the evolutionary process, these disabled connections serve as “pseudogenes” *in-vivo* that can reactivate in later generations due to mutation. Initially, the controller had 7 of them that, after pruning, we had 51 enabled connections to target. We used the same attribute to lesion the connections by virtually disabling them from passing values from source neurons to receivers. In other words, a lesion in our experiments means a severed connection in which, technically, would disrupt the flow of information from the source node to the receiver node. To lesion nodes, we then disabled the incoming/outgoing connections. For example, to lesion a neuron that sends information to three other neurons, we set those three connections to zero, which virtually silences the node.

Each lesion experiment started with silencing the targeted neuron or connection as described. All experiments consisted of 512 trials in which the network played the game 16 times per trial. The score of each trial was calculated by averaging these 16 scores, leading to a distribution of 512 scores per lesion experiment.

### Multi-perturbation Shapley Value Analysis

MSA is a rigorous method based on a Game-theoretical metric called Shapley value, here *γ* that indicates how much an element is important for the grand coalition. To elaborate, assume the marginal importance of an element *i* to a set of elements *S*, with *i* ∉ *S* is:

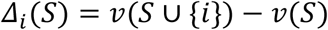

With *ν* being the worth or importance of the element *i*, and *S* a coalition of elements. Then *γ*_*i*_ while *i* ∈ *N* is defined as:

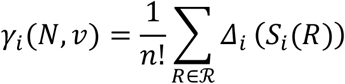

With ℛ is the set of all n! orderings of *N*, and *S*(*R*) is the set of elements preceding *i*, in the ordering *R*. We estimated *γ* of each neuron and connections by sampling 1000 orders from the permutation space of 19! for neurons and 51! for connections. These 1000 permutations then dictate which combinations of elements should be lesioned (Fig4). After selecting the target elements, we used the same perturbation approach as the single-site lesion and disabled the corresponding connections. The agent played the game 16 times, and the average score would be the score of that random permutation, providing a *γ* distribution of 1000 data points for each element. Altogether, we had around 70,000 unique combinations of lesions to estimate *γ* from.

### Statistical Inference

Besides testing the performance of the intact network against the random agent, blind, and weight-shuffled networks in which we used the non-parametric Mann-Whitney U test, we used bootstrap hypothesis testing to find significant statistics throughout the study. We first generated a synthetic null distribution for each statistical test by shifting the original distribution towards the H_0_’s mean value, either zero or an arbitrary number. For instance, to compare a distribution against a null distribution centered around zero, such as Shapley values, we subtracted the average from each data point, centered synthetic distributions around zero. In cases in which we tested distributions against a second distribution that is not centered around zero, such as the performance of the single-lesioned network versus the performance of the intact network, we shifted the synthetic distributions toward the H_0_’s mean, in this example, around 337 by adding the mean to each data point.

We then performed the bootstrapping and resampled the mean-adjusted distributions N times with replacement, with N being the number of original samples, e.g., 512 for single-site lesions. This generated a bootstrap dataset centered around the H_0_’s mean (Fig.12). We then calculated the bootstrap dataset’s mean and repeated the process 10,000 times to generate the bootstrap histogram of the means. In other words, the bootstrap histogram is a distribution of means if they were from a null hypothesis. We then checked if the mean values of our distributions fall above or below the p-value that is corrected for multiple comparisons using the Bonferroni correction method (0.05/Number of tests).

**Fig.12:**
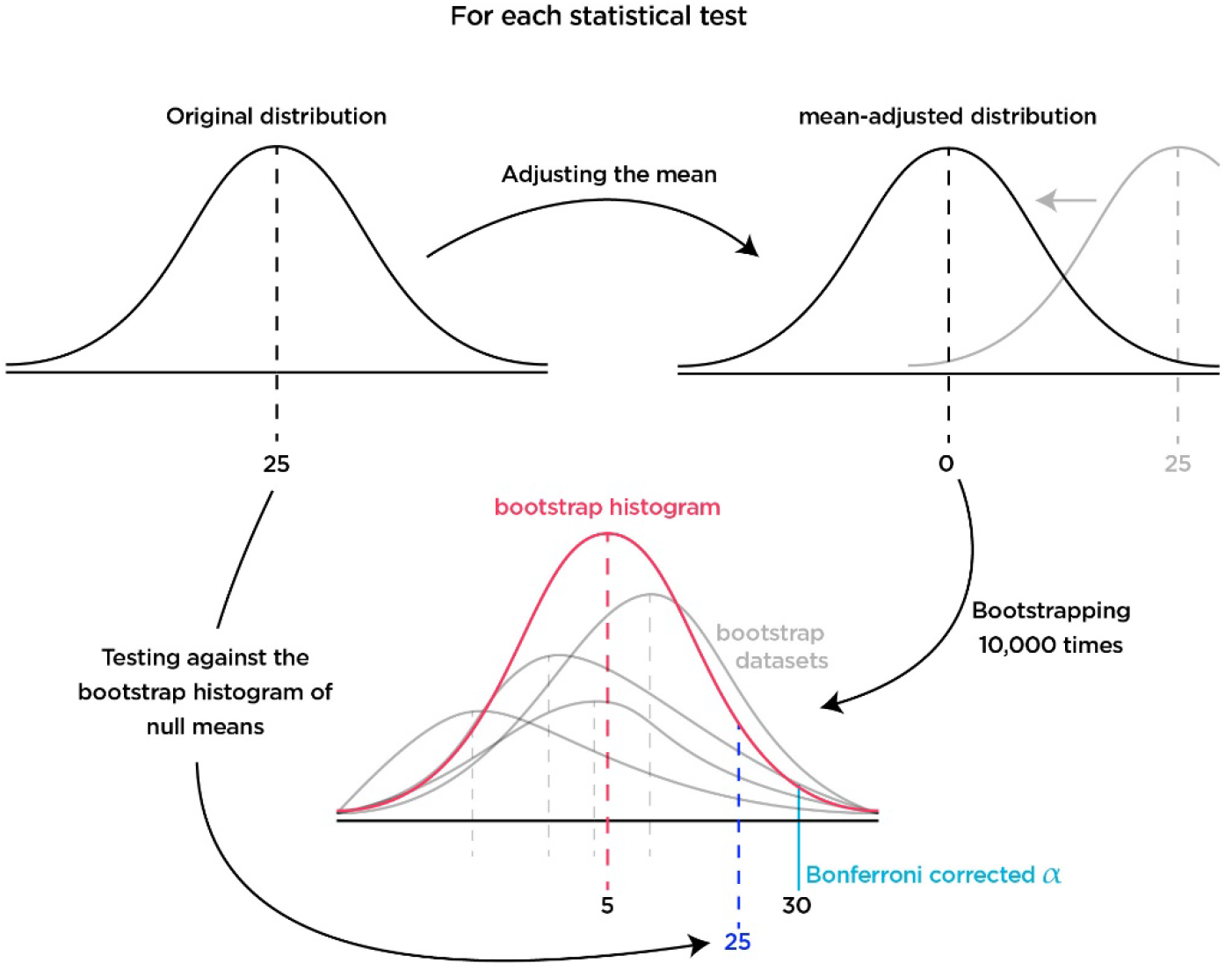
Visual diagram of the hypothesis testing process. For each test, we first made a null distribution by adjusting the mean. Then we resampled the synthetic distribution and kept track of the averages in the bootstrap histogram. Lastly, we checked if the original mean falls below or above the Bonferroni corrected p-value.

### Multivariate Transfer Entropy Analysis

We used the Information Dynamics Toolkit xl (IDTxl; [66]to analyze mTE between a set of targets (nodes 0 and 4) and sources (−1 and −4) in four conditions. First, the intact network, then the feedback loop from 0 to itself is lesioned, then the input from −4 to 0, and lastly, both the feedback loop and the input were lesioned. For each condition, we simulated 50 trials in which each trial had 1200 samples. We enforced this number by discarding trials with fewer samples and cutting the excessive samples from trials with more than 1200. Due to the quasi-binary dynamics of the target nodes, we used the Kraskov estimator instead of Granger causality to infer multivariate transfer entropy among the sources and targets. We further added information about the chosen action to the time series of the target nodes. If the node is chosen at time point t, then the value of the chosen node will be the value +1, and if not, just the raw data point (between 0 and 1) was stored. The reason was to account for saturation of the target nodes since, at some points, the actual values are very close to one another. Lastly, we injected a small amount of noise into the estimator (noise level = 1e-7). Both the minimum and maximum lag were set to 1 although we explored maximum lags of two and three. Eventually, we discarded the resulted lags and only reported the existence of TE between the pair of source and target since we found lags to be irrelevant for this analysis. To account for multiple comparisons, we set the number of omnibus permutations to 1000 and used the Bonferroni correction method to adjust the p-value (0.05/8), which sets the adjusted value to around 0.005.

## Supplementary Figures

**FigS1:**
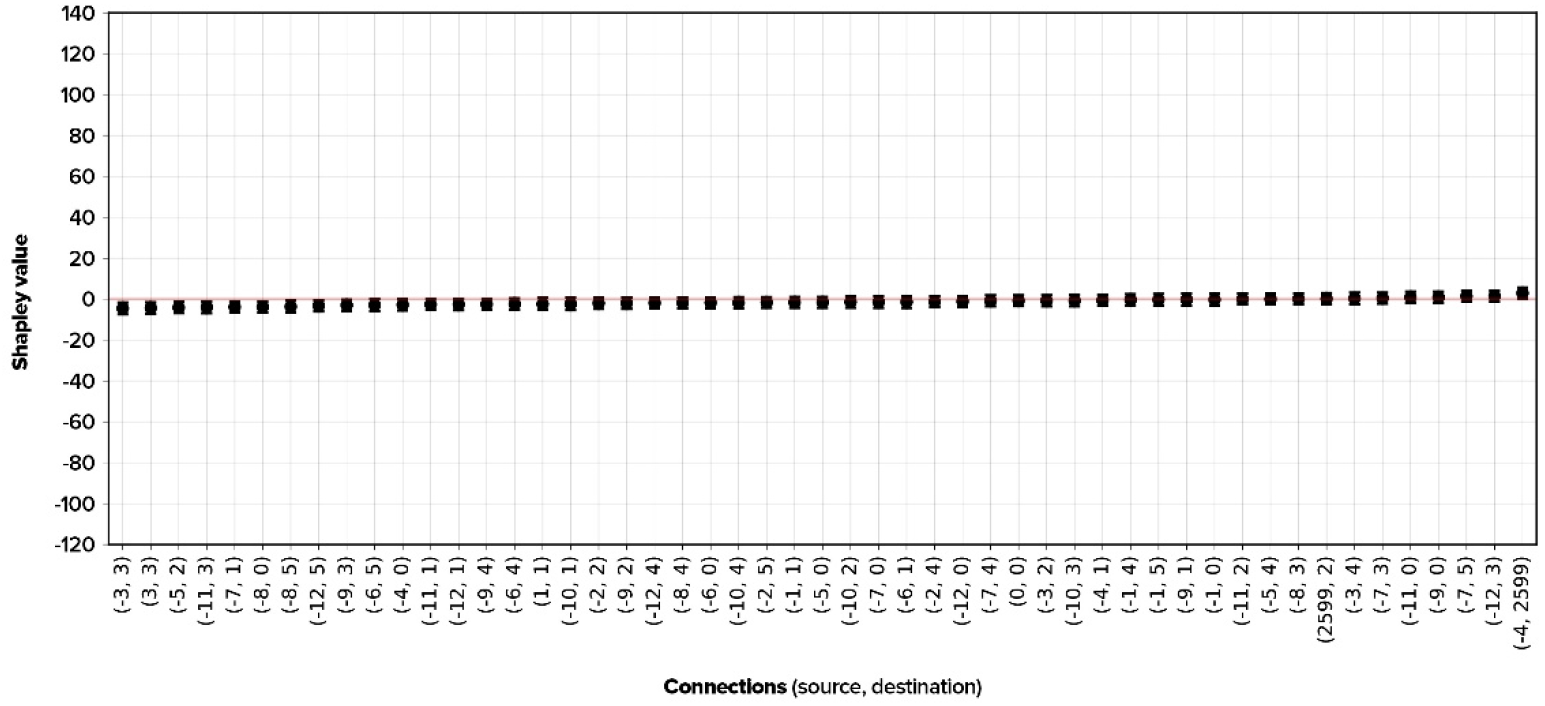
Shapley Values of the blinded network. As a sanity check, we performed the MSA on the optimized network connections while feeding it noise instead of game-states. The procedure is explained in the section: *Multi-perturbation Shapley value Analysis*. We found no connection with considerable causal importance since the network cannot perform properly.

## Acknowledgments

Funding is gratefully acknowledged: KF: Deutsche Forschungsgemeinschaft, Germany (SFB 936/A1; TRR169/A2), SMARTSTART, the joint training program in computational neuroscience of the Bernstein Network and the Volkswagen Foundation. CCH: Deutsche Forschungsgemeinschaft, Germany (SFB 936/A1; TRR169/A2; SFB 1461/A4; SPP 2041/HI 1286/7-1, HI 1286/6-1), the Human Brain Project, EU (SGA2, SGA3). Authors thank Alexandros Goulas, Fabrizio Damicelli, Fatemeh Hadaeghi, Joseph Lizier, Patricia Wollstadt, and Caroline Malherbe for their valuable comments and insights.

